# Transient cAMP production drives rapid and sustained spiking in brainstem parabrachial neurons to suppress feeding

**DOI:** 10.1101/2023.02.25.530033

**Authors:** Jonnathan Singh Alvarado, Andrew Lutas, Joseph C. Madara, Jeremiah Isaac, Caroline Lommer, Mark L. Andermann

## Abstract

Brief stimuli can trigger longer lasting brain states. G protein-coupled receptors (GPCRs) could help sustain such states by coupling slow-timescale molecular signals to neuronal excitability. Brainstem parabrachial nucleus glutamatergic neurons (PBN^Glut^) regulate sustained brain states such as pain, and express G_s_-coupled GPCRs that increase cAMP signaling. We asked whether cAMP directly influences PBN^Glut^ excitability and behavior. Both brief tail shocks and brief optogenetic stimulation of cAMP production in PBN^Glut^ neurons drove minutes-long suppression of feeding. This suppression matched the duration of prolonged elevations in cAMP, Protein Kinase A (PKA), and calcium activity *in vivo* and *in vitro.* Shortening this elevation in cAMP reduced the duration of feeding suppression following tail shocks. cAMP elevations in PBN^Glut^ neurons rapidly lead to sustained increases in action potential firing via PKA-dependent mechanisms. Thus, molecular signaling in PBN^Glut^ neurons helps prolong neural activity and behavioral states evoked by brief, salient bodily stimuli.

## Introduction

Behavioral states such as nausea, pain and associated arousal persist for minutes to hours. Such states are dependent on neural activity in the parabrachial nucleus^1–9^ (PBN) and are likely driven by slow-timescale changes in neural activity, yet how these persistent shifts in activity occur remains poorly understood. G_s_-coupled GPCRs signal via slow increases in cyclic AMP (cAMP). However, whether and how cAMP changes in any brain region control the duration of behavioral states has been challenging historically, due in part to technical limitations in studying cAMP signaling *in vitro* or *in vivo*. Recent advances have enabled the study of cAMP signaling in awake animals^10–14^. For example, in the case of mating drive, Gs-coupled GPCRs in the medial preoptic nucleus of the hypothalamus can convert brief neurotransmitter release into hours-long persistent behavioral states that match the duration of cAMP signaling^12^. However, depending on the set of regulators and downstream targets of cAMP signaling in a given cell type and brain area, cAMP signaling may persist for hours^12^ or only for seconds^11,15^ and may not necessarily couple directly to neural excitability^11,16^.

We focused on the PBN, a brainstem region where neurons express many GPCRs that drive or suppress production of cAMP^17,18^. We used new tools to selectively track and manipulate cAMP in PBN glutamatergic (PBN^Glut^) neurons. As the PBN is a key node in the control of ingestive, nausea, and pain-related behaviors, we investigated how cAMP signaling influences the duration of pain-evoked suppression of feeding, a behavior previously shown to involve the PBN^19^. We found that pain-evoked suppression of feeding lasts for tens of seconds, depends on sustained cAMP signaling, and can be mimicked by brief, light-evoked elevations in cAMP in PBN^Glut^ neurons. Using *in vivo* and *ex vivo* optical and electrophysiological approaches, we discovered that cAMP can rapidly control PBN^Glut^ neuron excitability. Thus, cAMP signaling provides a mechanism for translating various neuropeptidergic, hormonal and other signals acting on different PBN^Glut^ neurons into sustained neural activity and behavior, with additional specificity conferred by selective expression of GPCRs and phosphodiesterases that regulate the levels and dynamics cAMP in different PBN^Glut^ neuron types^18^.

## Results

### Brief salient events can halt feeding for extended periods of time

We first established a behavior that demonstrated sustained suppression of feeding for many tens of seconds following a brief, salient stimulus: shock-induced suppression of continuous feeding. We used a head-fixed behavioral setup because it allowed us to precisely deliver the same salient stimulus while measuring food consumption and could be easily combined with optical stimulation and recording methods (Figure 1A). Food restricted mice were acclimated to head restraint and trained to associate licking at a lickspout with the delivery of a palatable liquid food, Ensure (Figure 1B). Mice learned to lick vigorously at a steady rate, but stopped licking for an extended period of time (~30-60 seconds) following a 10-s tail shock stimulus (Figure 1B,C). Thus, despite a high overall motivation to consume food, the tail shock stimulus resulted in motivational suppression that outlasted the stimulus.

**Figure 1:**
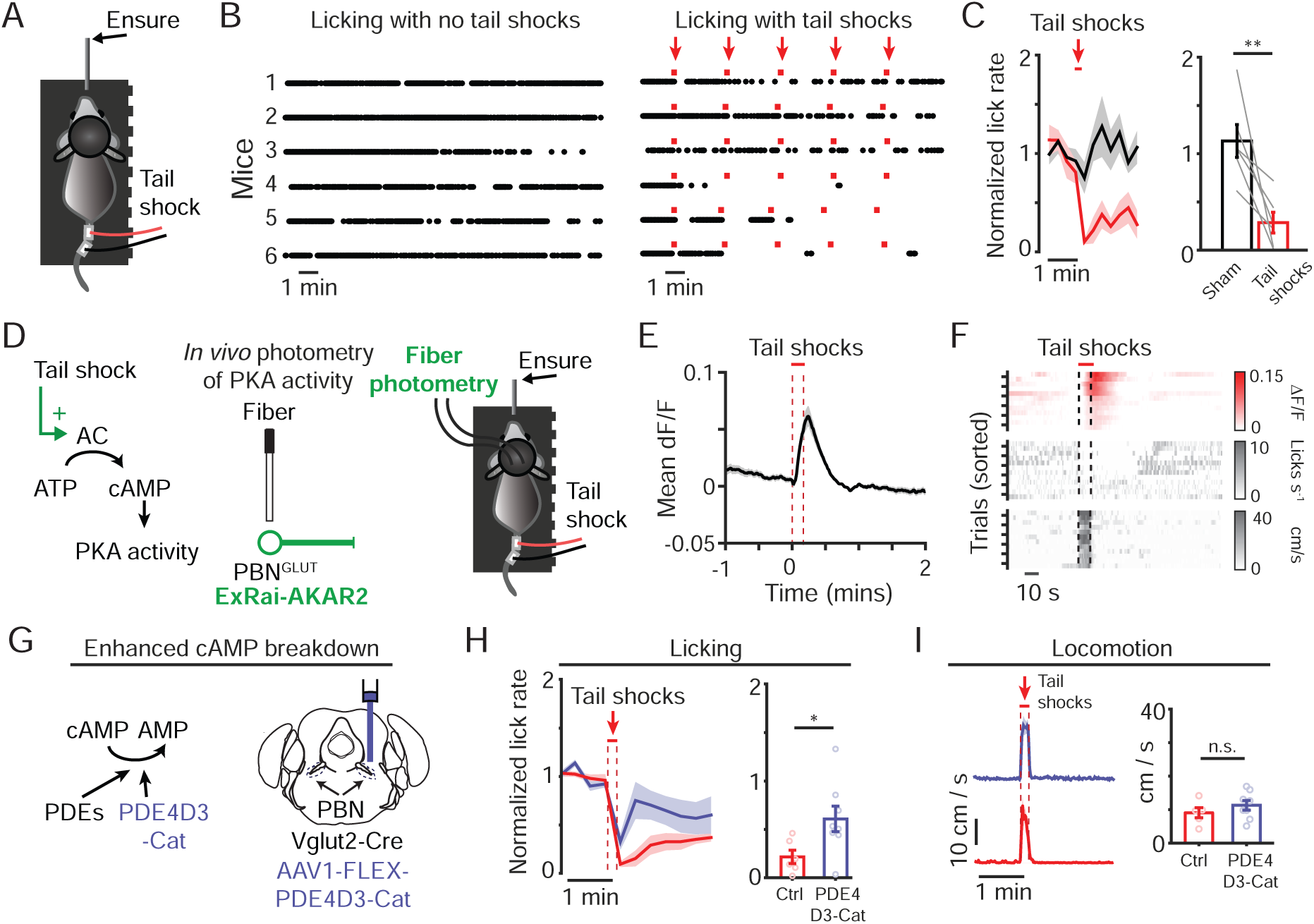
Brief, salient events halt feeding for extended periods of time via cAMP-dependent signaling in PBN^Glut^ neurons. A) Schematic depicting head-fixation on a spherical treadmill during consumption of liquid food (Ensure) and tail shock delivery. B) Left: licking during lick-triggered Ensure delivery on the control session in which no tail shocks were delivered. Right: same as left but on the test day with intermittent delivery of trains of tail shocks (10 s duration; 6 Hz; 0.35 mA). C) Left: time course of mean peak-normalized licking across mice (n = 6) on control (black; sham tail shock) vs. test sessions (red; tail shock delivery). Right: mean normalized licking in the two-minute period following tail shock delivery on the test sessions (red bar) or sham deliveries on control sessions (black bar). Student’s two-tailed paired *t*-test, p = 0.007. D) Left: schematic depicting predicted influence of tail shocks on adenylate cyclase activity in PBN neurons, leading to increased protein kinase A (PKA) activity. Middle: schematic depicting fiber photometry recording of a fluorescent reporter of PKA activity (ExRai-AKAR2). Right: schematic depicting fiber photometry recording during Ensure consumption and tail shock delivery. E) Time course of mean peak-normalized PKA activity aligned to the delivery of tail shocks (10 s; 6 Hz; 0.35 mA; n = 4 mice). F) Top: heatmap of normalized PKA activity (n = 12 trials from 4 mice) sorted by peak magnitude of PKA activity following tail shock delivery. Lick rasters (middle) and locomotion (bottom), plotted with same sorting as upper panel. G) Left: schematic depicting expression of the phosphodiesterase 4D3 catalytic domain (PDE4D3-Cat), which breaks down cAMP. Right: schematic depicting expression of PDE4D3-Cat unilaterally in PBN^Glut^ neurons. H) Left: time course of the mean normalized licking of Ensure and interruption by tail shocks in mice expressing PDE4D3-Cat (blue; n = 8 mice) or control mice (red; n = 6 mice). Right: mean normalized licking in the two-minute period following tail shock delivery in control mice (red bar) or mice expressing PDE4D3-Cat (blue bar). Student’s two-tailed unpaired *t*-test, p = 0.03. I) Left: time course of the mean locomotion (cm/s) for PDE4D3-Cat (blue) or control mice (red). Right: mean locomotion during the tail shock for mice expressing PDE4D3-Cat (blue bar) or control mice (red bar). Student’s two-tailed unpaired *t*-test, p = 0.42. For all panels, data are displayed as mean ± s.e.m. unless otherwise noted.

### cAMP-dependent signaling in PBN^Glut^ neurons controls prolonged suppression of feeding

We next asked whether molecular signaling in PBN^Glut^ neurons might underlie this extended duration of suppressed food consumption. Many subtypes of PBN^Glut^ neurons together play key roles in the processing of pain and in the suppression of food consumption^1,2,4–6,20,21^. Neuromodulators that signal through Gs-coupled GPCRs, which promote production of cAMP and activation of its main downstream effector, protein kinase A (PKA), have been shown to suppress feeding through their effects on PBN^20,22^. Thus, we asked whether PBN^Glut^ neurons exhibited prolonged PKA signaling driven by this transient tail shock, which could promote prolonged suppression of feeding following this brief experience (Figure 1D).

We took advantage of a genetically-encoded fluorescent reporter of PKA activity^11,12,23^ (exrai-AKAR2) which we previously found to be sensitive enough to detect tail shock-evoked PKA signaling^11^. We selectively expressed exrai-AKAR2 in PBN^Glut^ neurons and performed fiber photometry during shock-evoked feeding suppression (Figure 1D). We found that tail shock evoked an increase in PKA activity that peaked following shock offset and subsequently lasted for tens of seconds (Figure 1E). The evoked PKA signal was of variable amplitude but consistent duration across trials (Figure 1F). Accordingly, when we sorted individual trials by the magnitude of PKA activity following tail shock, we found that evoked PKA activity correlated with the duration of lick suppression (R = 0.82; p = 0.001), but not the duration of locomotion (R = 0.38; p = 0.22; Figure 1F). These results suggested that the strength and dynamics of cAMP-dependent signaling in PBN^Glut^ neurons may control PBN^Glut^ neuronal activity to arrest ongoing motivated behavior.

To directly test whether elevated cAMP in PBN^Glut^ neurons determines the duration of tail shock-evoked suppression of feeding, we expressed a constitutively active phosphodiesterase, which rapidly degrades cAMP, unilaterally in PBN^Glut^ neurons (Figure 1G; PDE4D3-Cat^11,12^). PDE4D3-Cat-expressing mice exhibited significantly shorter periods of feeding suppression following the tail shock, despite displaying a similar locomotor response during the tail shock (Figure 1H,I). The lack of effect on locomotion following disruption of cAMP signaling was consistent with our prior finding that PKA sensor activity correlated with licking, but not locomotion. Thus, intact cAMP accumulation in PBN^Glut^ neurons is necessary for the prolonged pause in feeding following tail shock.

### Transient photostimulated production of cAMP in PBN^Glut^ neurons suppresses food seeking for tens of seconds

Given the necessity of cAMP in setting the duration of feeding suppression following tail shock, we asked whether we could mimic this suppression by directly increasing cAMP in PBN^Glut^ neurons, thereby bypassing the transient activation of G_s_-GPCRs by diverse neuromodulators and peptides (Figure 2A,B). We expressed a blue-light activated adenylyl cyclase (biPAC^11,12^) bilaterally in PBN^Glut^ neurons to photostimulate cAMP production (Figure 2B). Photostimulation of cAMP production using low-intensity blue light (1 mW; 465 nm; five 1-s pulses spaced 2 s apart; Figure 2B) robustly suppressed ongoing food consumption and food seeking for tens of seconds following photostimulation (Figure 2C,D). Notably, this duration of feeding suppression was similar to that evoked by tail shock (Figure 1C).

**Figure 2:**
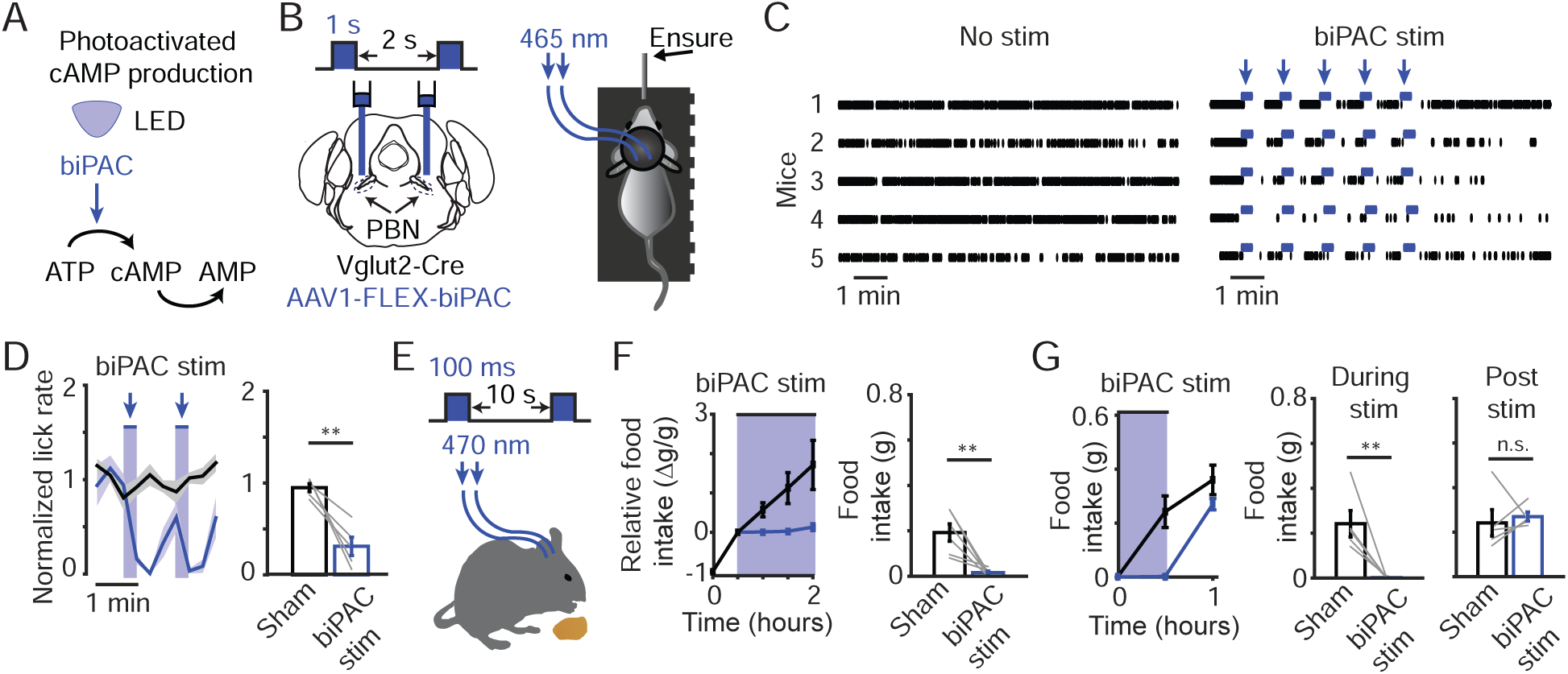
Transient photostimulated production of cAMP in PBN^Glut^ neurons suppresses food seeking for tens of seconds. A) Schematic depicting photostimulation of cAMP production via biPAC activation, and subsequent degradation of cAMP to AMP. B) Left: schematic depicting bilateral expression of biPAC in PBN glutamatergic neurons and photostimulation paradigm (1 s LED on, 2 s LED off; 10 s duration). Right: schematic depicting photostimulation of biPAC during Ensure consumption. C) Left: licking during lick-triggered Ensure delivery on the control session with no photostimulation of biPAC. Right: same as left but on the test session with intermittent photostimulation (465 nm; 1 mW). D) Left: time course of the mean normalized licking of mice (n = 5) on control (black; sham photostimulation) or test sessions (blue; photostimulation). Right: mean normalized licking in the one-minute period following photostimulation on test sessions (blue) or sham photostimulation on control sessions (black). Student’s two-tailed paired *t*-test, p = 0.006. E) Schematic depicting photostimulation of biPAC (100 ms LED on, 10 s LED off; 473 nm; 1 mW) during chow pellet consumption in freely moving mice. F) Left: food intake measurements over a two-hour session of refeeding in fasted mice without photostimulation (black; n = 6 mice) or with photostimulation (blue). Photostimulation started after the first 30 minutes. Right: mean food consumption during the last 90 minutes on control days without photostimulation vs. test days with photostimulation. Student’s two-tailed paired *t*-test, p = 0.009. G) Left: food intake measurements during the first hour of refeeding in fasted mice without photostimulation (black; n = 5 mice) or with photostimulation (blue; 1s LED on, 10 s LED off; 473 nm; 5 mW) during the first 30 minutes. Middle: mean food consumption during the first 30 minutes of sessions on control days without photostimulation vs. on test days with photostimulation. Student’s two-tailed paired *t*-test, p = 0.004. Right: mean food consumption during the first 30 minutes on control days without photostimulation vs. the 30 minutes following photostimulation on test days with photostimulation. Student’s two-tailed paired *t*-test, p = 0.67. For all panels, data are displayed as mean ± s.e.m unless otherwise noted.

Given the potency of these suppressive effects, we asked whether photostimulation of cAMP would also reduce home cage consumption of chow following a 24 hour fast. After allowing mice to consume food for 30 minutes, we bilaterally photostimulated cAMP production during the remaining 90 minutes of access to food using a low intensity stimulation paradigm (repeated cycles of 100 ms on; 10 s off; ~ 1 mW; 473 nm) that had no obvious effect on locomotor behavior (not shown) and that should maintain continuously elevated cAMP levels in PBN^Glut^ neurons (Figure 2E; see also Figure 3). Strikingly, we observed an almost complete absence of food consumption during this period (Figure 2F). In a separate experiment, we tested if suppression of feeding using a higher intensity biPAC photostimulation paradigm (repeated cycles of 1 s on; 10 s off; ~ 5 mW; 473 nm) would lead to sustained suppression of feeding long after the photostimulation ceased (Figure 2G). We found that mice resumed normal feeding following cessation of photostimulation, and that the total food consumption over the 30-minute period following cessation of photostimulation was similar to consumption during the initial 30-minute period of other sessions lacking photostimulation (Figure 2G).

**Figure 3:**
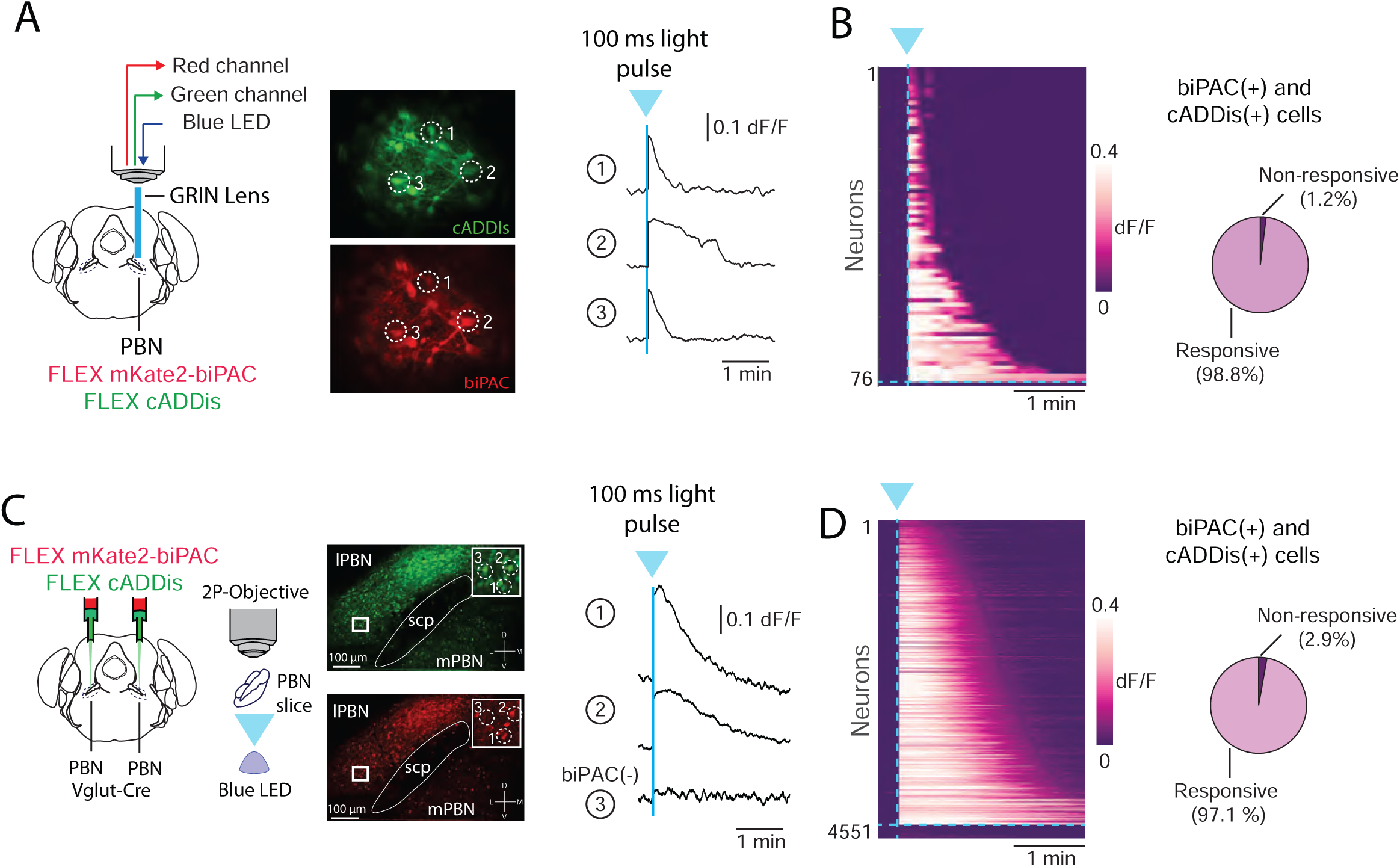
Photostimulated production of cAMP in PBN^Glut^ neurons persists for tens of seconds. A) Left: schematic depicting *in vivo* imaging approach for measuring and stimulating cAMP using cADDis and biPAC. Right: representative traces of cADDis fluorescence from 3 neurons stimulated with blue light (brief 100 ms pulse). B) Heat map showing effect of biPAC stimulation on all imaged neurons (76 neurons, 4 mice). Right: fraction of neurons that displayed cADDis responses to blue light stimulation. C) Left: similar to (A), but imaging in brain slices. Right: representative traces of cADDis fluorescence from 3 neurons stimulated with blue light (100 ms pulse). D) Heat map showing effect of biPAC stimulation on all neurons (4551 neurons, 15 slices from 5 mice). Right: fraction of neurons that displayed cADDis responses to blue light stimulation.

### Photostimulated production of cAMP neurons persists for tens of seconds in some PBN^Glut^ neurons

Having mimicked the sustained time course of shock-evoked suppression of feeding using brief biPAC photostimulation of cAMP, we next sought to directly assess the dynamics by which cAMP in individual PBN neurons decays to baseline following photostimulation. We co-expressed the fluorescent cAMP biosensor cADDis^10–12^ and biPAC in PBN^Glut^ neurons, implanted a gradient index (GRIN) lens in lateral PBN, and performed two-photon imaging in awake mice (Figure 3A). Brief photostimulation of cAMP using the same stimulation duration as in Figure 2F (100 ms) evoked robust increases in cAMP in all PBN^Glut^ neurons expressing both biPAC and cADDis (Figure 3B). The elevation in cAMP persisted for tens of seconds following photostimulation before decaying back to baseline (median *τ*, 33.2 seconds). These rapid decay dynamics^11^ indicate that PBN^Glut^ neurons contain abundant levels of endogenous phosphodiesterases that break down cAMP into AMP, in contrast to the slower dynamics observed in the medial preoptic nucleus of the hypothalamus using a similar protocol^12^.

The GRIN lens imaging was restricted to a local subregion of lateral PBN (effective FOV: ~250 μm^2^). To investigate whether similar signaling dynamics exist across all regions of PBN, we repeated the photostimulation protocol in coronal brain slices, which allowed imaging of hundreds of neurons across lateral and medial PBN (Figure 3C). Similar to the above results, we found that all imaged PBN neurons showed elevated and sustained cAMP following biPAC photostimulation. Given the diversity of cell types in the PBN^18^, we characterized the decay dynamics across PBN neurons, as this could provide a means by which distinct PBN neurons differentially integrate peptidergic and neuromodulatory inputs across fast or slow time scales. To this end, we plotted the time constants for individual PBN neurons in all FOVs (Supplemental Figure 1A). The decay time constants of cAMP clearance were highly variable between neurons both within and across brain slices, varying between ~10 and several hundred seconds, but were not systematically different on average across the major subdivisions of PBN (Supplemental Figure 1A-C).

### Photostimulation of cAMP drives persistent increases in calcium activity in some PBN^Glut^ neurons

Depending on the downstream molecular machinery in any given neuron, elevations in cAMP may or may not modify neural activity on a rapid timescale^11,12,16^. Given the correspondence between elevated cAMP levels in PBN^Glut^ neurons and feeding suppression, we hypothesized that increases in cAMP lead to immediate and sustained increases in neuronal activity in at least some PBN^Glut^ neurons. To test this prediction, we performed *in vivo* recordings of neuronal calcium signals as a proxy for spiking in PBN^Glut^ neurons co-expressing GCaMP6s and biPAC. We focused on neurons with confirmed biPAC expression and rapid GCaMP6s fluctuations (Figure 4A). Photostimulated cAMP production evoked GCaMP6s fluorescence transients in a subset (34%) of PBN^Glut^ neurons, suggesting that some but not all can translate cAMP into neuronal activity (Figure 4B). Notably, these GCaMP responses to the 100 ms light pulse could last for several tens of seconds. While biPAC expression levels were correlated with the amplitude of photogenerated cAMP, they were not correlated with the magnitude of light-evoked GCaMP activity (Supplemental Figure 1D-I).

**Figure 4:**
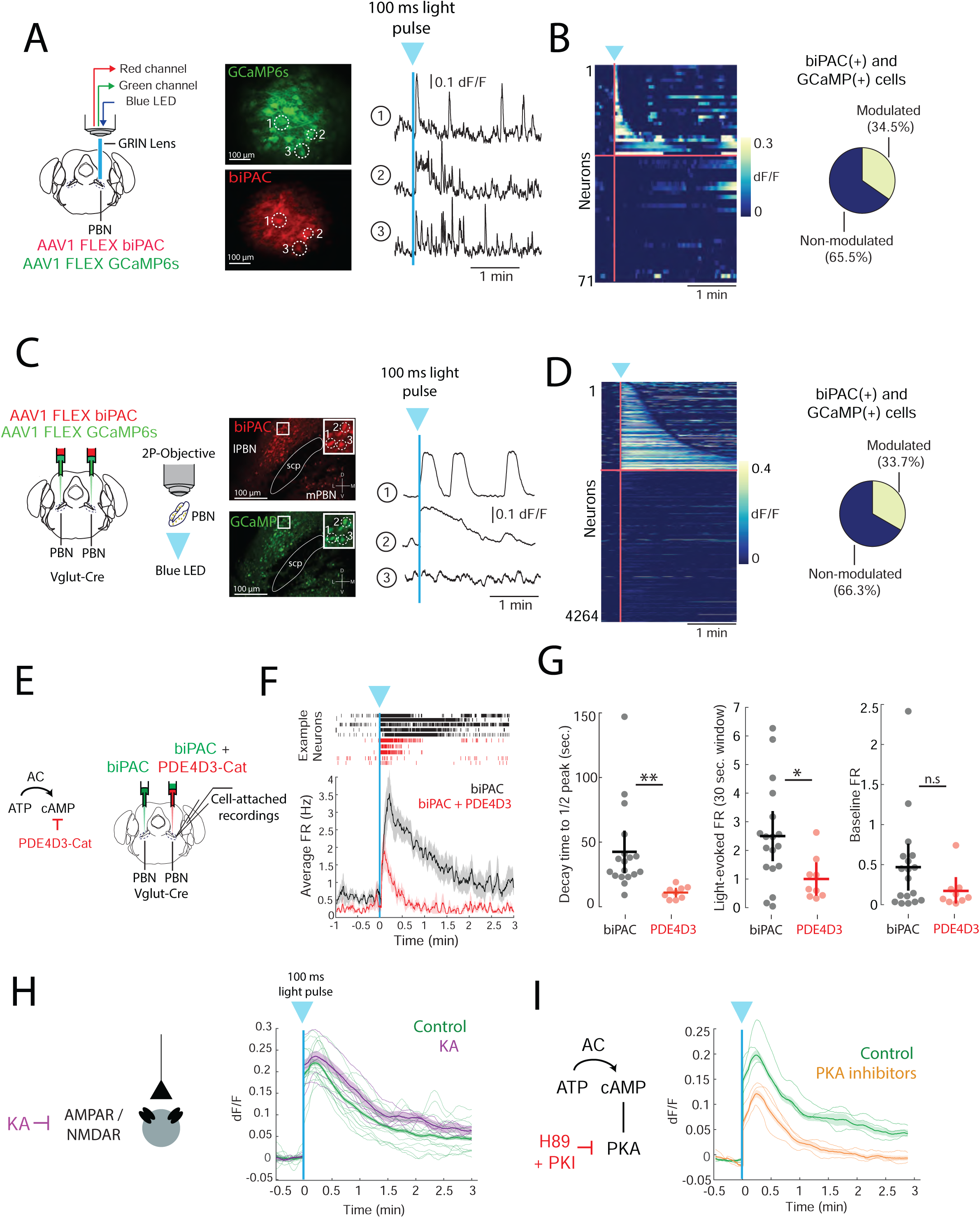
Photostimulation of cAMP drives persistent increases in calcium activity in some PBN^Glut^ neurons. A) Left: schematic depicting *in vivo* imaging approach for measuring calcium activity using GCaMP and stimulating cAMP using biPAC. Right: representative traces of GCaMP fluorescence from 3 neurons stimulated with blue light (100 ms pulse). B) Heat map showing effect of biPAC stimulation on all neurons (71 neurons, 4 mice). Right: fraction of neurons that displayed GCaMP responses to blue light stimulation. C) Left: similar to (A), but imaging in brain slices. Right: representative traces of GCaMP fluorescence from 3 neurons stimulated with blue light (100 ms pulse). D) Heat map showing effect of biPAC stimulation on all neurons (3330 neurons, 17 slices from 5 mice). Right: fraction of neurons that displayed GCaMP responses to blue light stimulation. E) Schematic of viral infection strategy for cell-attached recordings in PBN. F) Top: example neurons (5/condition) showing single-trial firing rate responses to blue light in the presence (red) or absence (black) of PDE4D3-Cat expression. Bottom: average firing rate for all recorded neurons in each condition. G) From left to right: group data showing time to half peak (\*\**p* < 0.01, *t* = 2.82, Student’s two-tailed unpaired *t*-test), mean firing rate response (*p < 0.05, t = 2.42, Student’s two-tailed unpaired *t*-test), and mean baseline firing rate for control and PDE4D3-Cat conditions (p = 0.16, Student’s two-tailed unpaired *t*-test). Cells expressing PDE4D3-Cat also displayed slightly reduced firing rates in the pre-stimulation window, though this result did not reach statistical significance. H) Illustration of the effect of kynurenic acid (KA) on excitatory synaptic transmission. Right: average GCaMP responses to biPAC stimulation in the presence (381 responsive neurons, 4 slices from 3 mice) or absence of KA (1702 responsive neurons, 17 slices from 7 mice). p = 0.10, Student’s two-tailed unpaired *t*-test. I) Illustration of the effect of PKA inhibitors. Right: average GCaMP responses in the same neurons before and after application of PKA inhibitors (411 responsive neurons, 4 slices from 3 mice). Mixed effects model, drug effect size: 0.075 dF/F, *** p < 0.001). Mean dF/F is calculated over the 30 seconds following stimulation. Data are displayed as mean ± s.e.m.

To study the mechanism(s) underlying this cAMP-driven PBN^Glut^ neuron activity, we again turned to acute brain slices. Upon photostimulation, ~32% (1437/4264) of neurons displayed a rapid increase in fluorescence, similar to the proportion observed *in vivo* (Figure 4C-D). We identified two categories of responsive neurons: those with a transient light-evoked elevation in calcium that seemed to mirror the cAMP transient (~53% of responsive neurons; median duration 38.2 s), and those that displayed a more sustained elevation in neural activity that outlasted the typical duration of cAMP (~47% of responsive neurons, Supplemental Figure 2A,B; median duration 121.6 s). Roughly similar proportions of cAMP-responsive neurons with either response profile were observed in each slice. Notably, the duration of GCaMP responses tended to be longer in the dorsomedial part of lPBN, and briefer in the ventrolateral part (Supplemental Figure 2C,D), despite a lack of systematic spatial differences in the duration of evoked cAMP responses across these two regions of lateral PBN (Supplemental Figure 1A-C).

The above findings indicate that most PBN^Glut^ neurons show prolonged elevations in cAMP following biPAC stimulation, while a smaller subset also shows increased calcium activity. We wanted to directly test whether the capacity to directly convert cAMP elevations into increases in calcium activity is indeed restricted to a subset of PBN^Glut^ neurons, or whether these results were due to our biPAC protocol for elevating cAMP. To this end, we also applied the adenylyl cyclase activator forskolin (20 μM) while simultaneously recording both the green cAMP sensor, cADDis, and the red-shifted calcium indicator, jRGECO. Consistent with the above findings during biPAC stimulation, the majority of PBN^Glut^ neurons expressing cADDis (92%) first showed increased cAMP upon forskolin application, while only a fraction of these neurons (~44%) showed a subsequent increase in calcium activity (Supplemental Figure 3A-C). Given that some of the observed increases in calcium signals could reflect subthreshold changes in calcium, we next examined the relationship between cAMP-induced calcium activity and spiking activity in greater detail.

### cAMP elevates action potential frequency in PBN^Glut^ neurons

We first confirmed that the increases in GCaMP6 fluorescence following brief biPAC-evoked stimulation of cAMP reflected sustained increases in spiking activity of PBN^Glut^ neurons, as they were largely absent when we blocked the ability of neurons to spike using tetrodotoxin (TTX, 2 μM, Supplemental Figure 4A,B). To further assess the magnitude and dynamics of this increase in spiking, we performed cell-attached electrophysiological recordings of PBN^Glut^ neurons expressing biPAC in brain slices. Photostimulation of cAMP generated a potent increase in firing rate that peaked ~5 seconds after stimulation and slowly decayed to baseline (Figure 4E,F), approximately matching the time course of our *ex vivo* GCaMP6 imaging experiments. In contrast to cell-attached recordings, this firing rate increase was not observed during whole-cell recordings (data not shown), suggesting that intact intracellular signaling is necessary for the translation of cAMP to spiking activity in PBN^Glut^ neurons.

A key finding in Figure 1 was that the duration of tail-shock-evoked feeding suppression was blunted by expression of a constitutively active phosphodiesterase, PDE4D3-Cat. To directly confirm that PDE4D3-Cat expression curtails the duration of firing rate increases, we performed cell-attached recordings in PBN^Glut^ neurons that expressed either biPAC alone, or a combination of biPAC and PDE4D3-Cat (Figure 4E). Spiking responses to photostimulation of the biPAC-only hemispheres were similar to bilateral biPAC-only recordings (and thus we combined both datasets). In contrast, PBN^Glut^ neurons co-expressing biPAC and PDE4D3-Cat showed weaker increases in firing rate in response to photostimulation (Figure 4F,G; biPAC only: 2.45 ± 0.4 Hz; biPAC and PDE4D3-Cat: 0.95 ± 0.3 Hz) and decayed more quickly to baseline spiking levels (decay time to half-peak, biPAC only: 42.11 ± 7.7 s; biPAC and PDE4D3-Cat, 10.6 ± 2.4 s).

### PKA, but not excitatory synaptic transmission or HCN channels, is required for the coupling between cAMP and neuronal activity

To better understand how cAMP is translated into neuronal activity, we performed a series of pharmacological experiments in our GCaMP slice imaging preparation. To test whether cAMP increased neural activity via increased synaptic transmission, we stimulated biPAC in the presence of an antagonist of glutamate receptors, kynurenic acid (KA, 2mM). Both the transient and sustained phases of neuronal activity induced by biPAC stimulation were largely unchanged in the presence of KA (Figure 4H).

cAMP can influence neuronal activity through downstream effectors such as PKA, as well as via regulation of hyperpolarization-activated cyclic nucleotide gated (HCN) channels^26,27^. We therefore sought to test the contribution of these downstream pathways to biPAC-evoked activity. Application of a selective blocker of HCN channels (50 μM ZD7288) did not significantly influence biPAC-evoked activity (Supplemental Figure 4C,D). In contrast, PKA inhibitors (800 nM PKI and 10 μM H-89) substantially attenuated biPAC-evoked activity (Figure 4I, Supplemental Figure 4E,F), indicating a significant role for this pathway in translating cAMP into neuronal activity.

## Discussion

Through the activation of cAMP and other second messengers, GPCRs can signal beyond the duration of ligand binding. Here, we studied how cAMP influences neural activity within the PBN, a brain region that processes multimodal bodily signals to influence behaviors over multiple timescales and expresses a wide variety of GPCRs^18^. Given how little is known about cAMP signaling in PBN, we chose to investigate all PBN glutamatergic neurons, which account for ~80% of all PBN neurons, as the results of this study could be of value across the multiple molecularly defined PBN cell types. Using a suite of tools to monitor and manipulate cAMP *in vivo* and *ex vivo*, we show that photostimulated and pharmacologically evoked elevations in cAMP persist for tens of seconds to minutes in PBN^Glut^ neurons before returning to baseline levels. The sustained elevation in cAMP and downstream PKA signaling in PBN^Glut^ neurons was necessary for sustained suppression of feeding after tail shock, as expression of PDE4D3-Cat, which attenuates cAMP accumulation, also abbreviated shock-evoked suppression of feeding. Direct optogenetic induction of cAMP production was also sufficient to phenocopy the prolonged shock-evoked suppression of feeding. By directly controlling production of cAMP, we found that sustained increases in cAMP are directly translated into similarly sustained increases in neural activity in roughly one third of PBN^Glut^ neurons via PKA-dependent mechanisms. Thus, in contrast to the more common roles of cAMP as a slow regulator of synaptic plasticity, development, and learning^15,28^, our findings demonstrate a more direct role for cAMP in increasing neuronal spiking in PBN. Specifically, in response to brief, salient bodily signals, cAMP can drive sustained PBN neuron activation and associated suppression of appetitive behaviors on timescales from tens of seconds to minutes.

### Endogenous neuromodulatory and peptidergic drivers of cAMP in PBN neurons

Our photometry experiments were designed to investigate the contribution of prolonged cAMP signaling to the sustained behavioral response to a brief, salient stimulus (tail shock). An outstanding question is which endogenous ligands and receptors drive relevant cAMP signaling in PBN neurons. Indeed, previous work has identified unique patterns of GPCR expression across PBN cell types^18^. One potentially important neurotransmitter is dopamine, which recently was shown to be causally involved in the control of feeding and body weight by PBN neurons^22^. Both cAMP-producing Drd1 and cAMP-inhibiting Drd2 receptors are expressed in PBN neurons, and dopamine-expressing neurons in multiple brain areas project to the PBN (e.g., from VTA, DRN, NTS)^20,29,30^. Another important neuropeptide signal in the PBN is neuropeptide-Y (NPY), which via NPY1/2R leads to decreased cAMP and has been demonstrated to counter the ability of painful stimuli to suppress feeding^7^. In addition, the mu opioid receptor, another GPCR that drives reductions in cAMP, is highly expressed in the PBN and plays important roles in regulating breathing and pain sensitivity^31^. Future studies will be necessary to understand whether and how GPCR expression patterns contribute to sensory processing in PBN. Given the direct coupling of cAMP to spiking activity in many PBN^Glut^ neurons, it is likely that variable expression of different Gs-coupled (cAMP-producing) GPCRs across distinct PBN neuron subtypes allows for specific neuromodulatory/peptidergic contexts that facilitate and/or mimic the relay of specific bodily signals to the rest of the brain.

### Determinants of the duration of *in vivo* cAMP signaling in PBN

Recent work has revealed that neuromodulator release can be integrated over long periods of time (minutes to hours) via cAMP. For example, release of dopamine from hypothalamic dopamine neurons upon a brief encounter with a female drives hour-long increases in cAMP in medial preoptic neurons in male mice, which in turn causes a priming of mating behaviors that persists for a similarly long period of time^12^. Related work in fruit flies shows that cAMP signaling through PKA serves as a persistent biochemical signal even in the absence of continuous synaptic input, allowing for cell-autonomous persistent activity^16^. Similar processes might take place in the PBN, where cells responsive to acute “threats”^5,32^ (pain, noxious temperatures, toxin consumption) trigger defensive behavioral responses that can persist beyond the duration of the initial threat stimulus^4,32^. Future studies that leverage cAMP sensors with increasing signal-to-noise ratios and insensitivity to bleaching^33,34^ will be ideally situated to address these questions.

The duration of elevated cAMP (and the consequent effects on spiking and behavioral suppression) may be modifiable via regulation of endogenous phosphodiesterase activity in PBN^Glut^ neurons. The phosphosdiesterases PDE4 and PDE1 are highly expressed in PBN^35^ and PDE1 is activated by calcium^36^, indicating a possible negative-feedback mechanism by which neural activity can shorten the duration of future elevations in cAMP. Future studies can investigate how changes in phosphosdiesterase activity or expression levels across minutes to days, and across subtypes of PBN^Glut^ neurons, may impact the persistence of cAMP and spiking in PBN.

### Coupling of cAMP to neuronal activity

We showed that while most neurons in PBN had the capacity for sustained cAMP signaling across tens of seconds to minutes, photostimulation of cAMP production drove spiking-related activity in only a subset (~one third) of PBN^Glut^ neurons. What determines which neurons in PBN ultimately translate cAMP into neural activity? One possibility is that differential expression of PKA, a canonical target of cAMP, determines the specificity of each neuron’s response. The mechanism by which PKA increases excitability remains to be determined, but could include phosphorylation of ion channels that are differentially expressed across subtypes of PBN^Glut^ neurons^18^. Moreover, most subunits of PKA appear highly expressed throughout the entire PBN (Allen Brain Atlas), but distinct isoforms appear biased towards different PBN subregions. This differential expression could account for the spatial gradient we observed, where activity driven by cAMP is more sustained in dorsomedial lPBN, and relatively more transient in ventrolateral lPBN. Notably, our experiments were carried out in PBN^Glut^ neurons (which make up ~80% of all PBN neurons^2,9^) suggesting that coupling of cAMP to sustained spiking is a relatively common feature in PBN. Future experiments could aim to better understand how this feature impacts more specific interoceptive modalities controlled by distinct PBN cell types^18^.

The observation that PKA inhibitors only partially abolished biPAC-evoked spiking suggests that PKA-independent mechanisms could also contribute to the coupling of cAMP to spiking. Cyclic nucleotide-gated (CNG) channels and exchange proteins directly activated by cAMP (EPAC) can directly depolarize neurons^37,38^, allowing for PKA-independent regulation of spiking. Thus, while we ruled out cAMP-activated HCN channels as the main relevant channel, future work could more precisely define the mechanism(s) by which cAMP regulates neuronal excitability in lPBN.

### Potential limitations of this study

Here, we have focused on investigating the role of cAMP signaling in the PBN on acute suppression of feeding. To observe the effect of cAMP in an unbiased manner, we extended our focus to all PBN^Glut^ neurons, revealing heterogeneous timescales for cAMP and cAMP-driven activity across all of PBN. Thus, more studies will be needed to investigate how cAMP influences the activity of distinct PBN cell types as well the timing of their associated behaviors such as control of respiration^31,39,40^, malaise-induce anorexia^1,4,41,42^, thermoregulation^43,44^, feeding^22,45–49^, and pain sensitization^2,9, 50–53^. Our ability to monitor cAMP dynamics during natural behaviors was hampered by the sensitivity of the cADDis sensor *in vivo*, which motivated us to use other sensors such as exrai-AKAR2 to track PKA activity as a proxy for cAMP. Newly emerging tools may hopefully allow for even finer interrogation of cAMP signaling^23,33,34^, particularly on longer timescales (hours to days), and during more naturalistic conditions.

## Acknowledgements

We thank B. Lowell, M. Krashes, S. Liu, S. Zhang, R. Essner, K. Ruda and other members of the Andermann laboratory for useful feedback. We also thank S. Zhang for technical assistance with biPAC stimulation experiments. Authors were supported by NIH F32 DK112589, a Davis Family Foundation award, a BNORC pilot grant (P30DK046200), and ZIA DK075169 (A.L.); a Harvard Mind Brain and Behavior postdoctoral fellowship (J.S.A), NIH DP2 DK105570, R01 DK109930, DP1 AT010971, R01 MH12343, a McKnight Scholar Award, and grants from the Boston Nutrition and Obesity Research Center (P30 DK046200) and the Klarman Family Foundation (M.L.A.); and the Boston Children’s Hospital Viral Core for virus packaging (NIH P30 EY012196).

## Author Contributions

Conceptualization, J.S.A., A.L. and M.L.A.; investigation, A.L, J.S.A., J.M., J.I., and C.L.; formal analysis & visualization, A.L., J.S.A., and M.L.A.; writing – original draft, J.S.A. and A.L., writing – review & editing, J.S.A., A.L., and M.L.A.

---

**Supplemental Figure 1.**
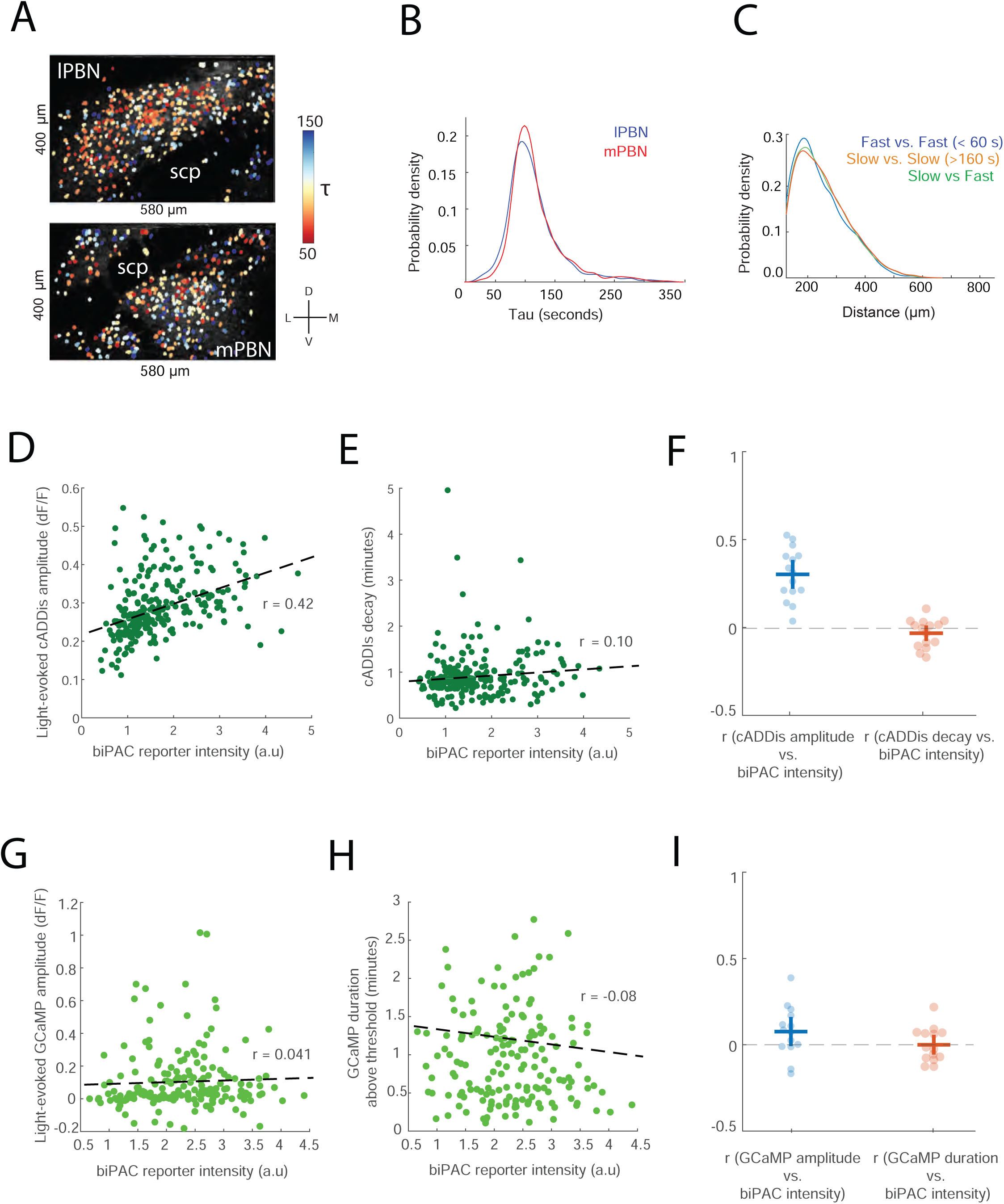
Relationship between biPAC expression levels and evoked cAMP and neuronal activity. Related to Figure 3. A) ROIs for PBN^Glut^ neurons coexpressing cADDIs and biPAC colored by the estimated decay time constant *τ* in two example slices. B) Histogram of *τ* values for neurons located in lateral PBN (lPBN; 3843 neurons) and medial PBN (mPBN; 532 neurons). C) Histograms of distances between pairs of fast-decay neurons, pairs of slow-decay neurons, or between pairs including one fast-decay and one slow-decay neuron. D) Scatter plot of biPAC reporter intensity (a rough measure of biPAC expression level) vs. cADDis response magnitude (average over first 20 seconds) to blue light for one representative slice. R: correlation coefficient computed for this example slice. E) Scatter plot of biPAC reporter intensity vs. cADDis signal decay rate for one representative slice. F) Group data showing correlation coefficients for each slice for analyses in (D) and (E) (15 slices, 6 mice). G) Scatter plot of biPAC reporter intensity vs. GCaMP response magnitude (average over first 20 seconds) to blue light for one representative slice. R: correlation coefficient. H) Scatter plot of biPAC reporter intensity vs. GCaMP duration above threshold. I) Group data showing correlation coefficients for each slice for analyses in (G) and (H) (14 slices, 5 mice). Data are displayed as mean ±s.e.m.

**Supplemental Figure 2.**
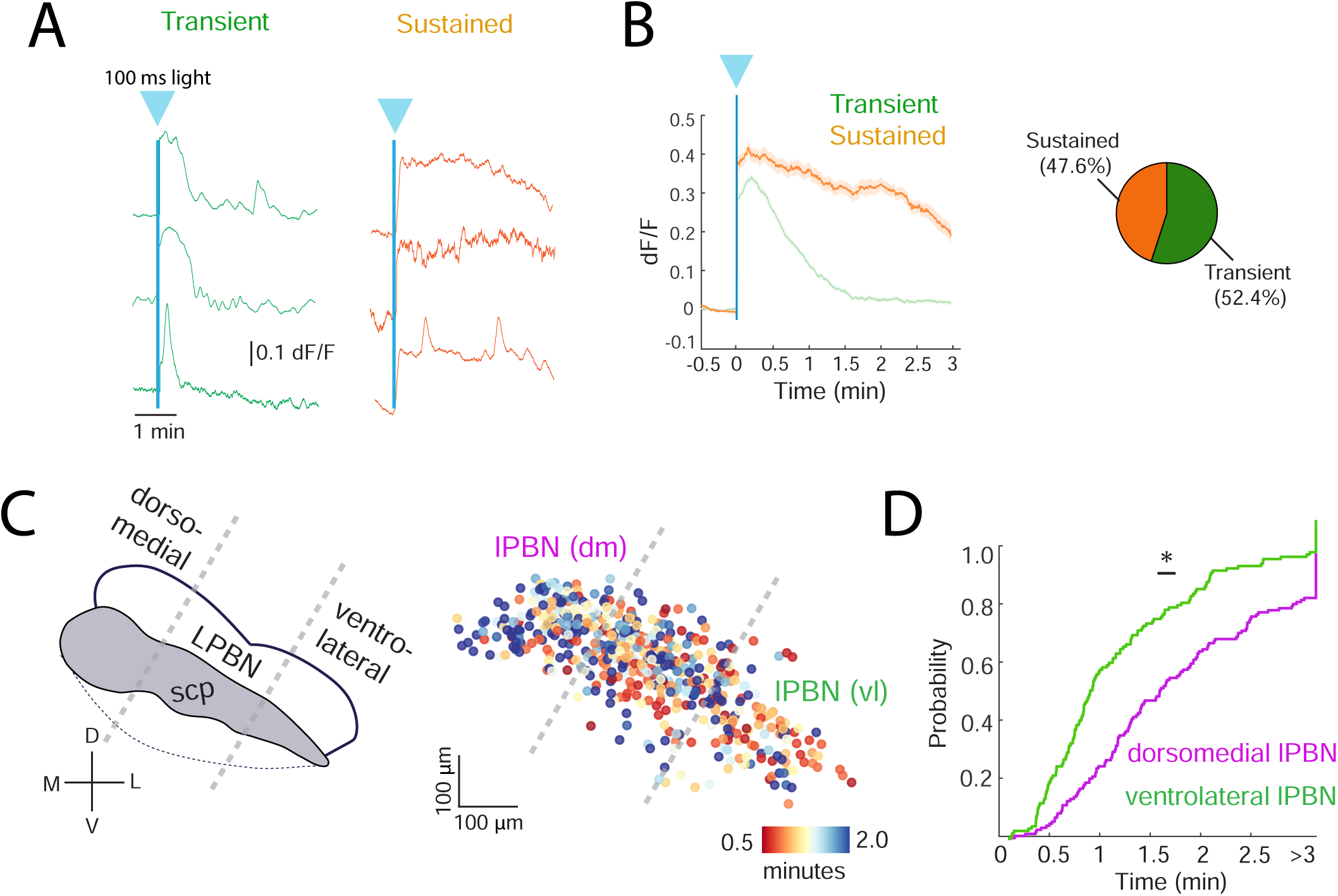
Timescale and spatial organization of responsive GCaMP-expressing neurons. Related to Figure 4. A) Example transient (left) or sustained (right) GCaMP responses evoked by 100 ms of biPAC stimulation (blue arrow and line) in one slice imaging experiment. B) Average GCaMP time courses for all transient (n = 509) and sustained (n = 371) neurons (15 slices from 6 mice). Right: proportion of biPAC-responsive neurons that were classified as transient or sustained. C) Left: diagram of two major subdivisions of PBN that were considered for spatial analyses. Right: map of neurons in PBN, colored by GCaMP response duration (time above threshold, see Methods), from 6 lPBN FOVs that had more than 50 responsive neurons. lPBN was subdivided into three sections along the dorsomedial to ventrolateral axis. The two flanking sections, ventrolateral (vl) and dorsomedial (dm) lPBN, were used for quantification. The middle section was ignored to minimize the effect of alignment error in subsequent analyses. D) Cumulative distribution function of GCaMP response durations for ventrolateral and dorsomedial lPBN.

**Supplemental Figure 3.**
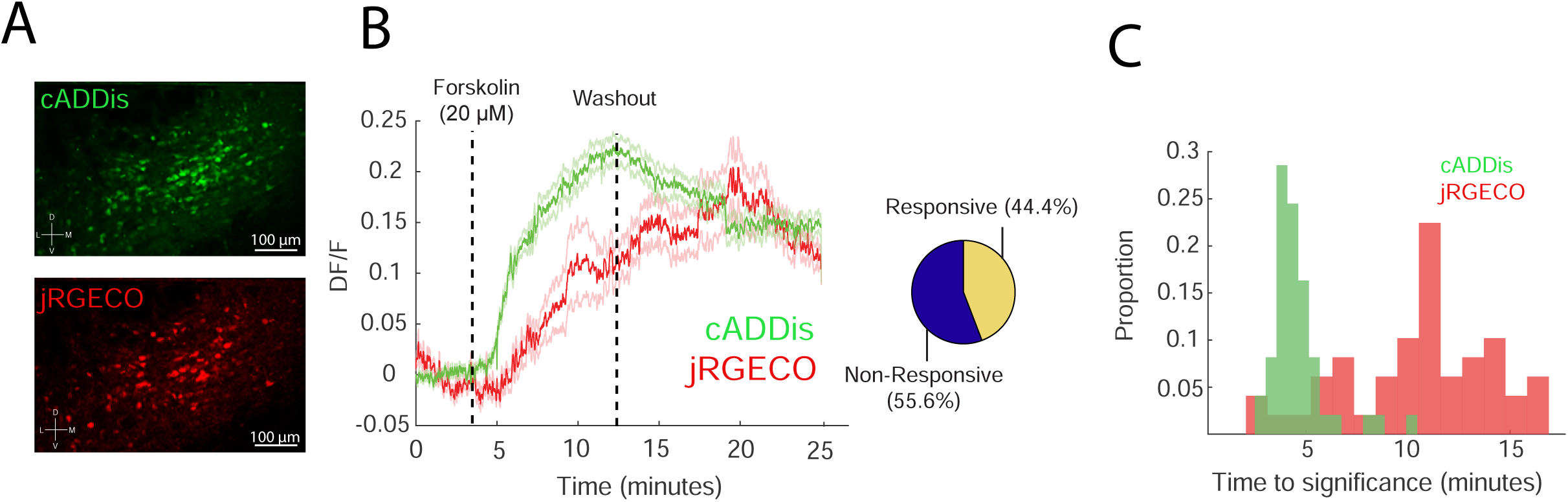
Effect of pharmacologically driven cAMP production on neuronal activity. Related to Figure 4. A) Representative slice imaging experiment showing expression of jRGECO1 and cADDis. B) Average time course of cADDis and jRGECO1 activity during application and washout of forskolin. Right: proportion of neurons that responded to forskolin application (84 responders out of 190 neurons that expressed both jRGECO1 and cADDIs; 2 slices from 2 mice). Note that cADDis is a downward sensor, and therefore increases in cAMP and calcium involve changes in fluorescence in opposite directions. C) Latency for cADDis or jRGECO1 to reach significant activation after application of forskolin (median duration for cADDis, 2.7 minutes; median duration for jRGECO1, 9.5 minutes).

**Supplemental Figure 4.**
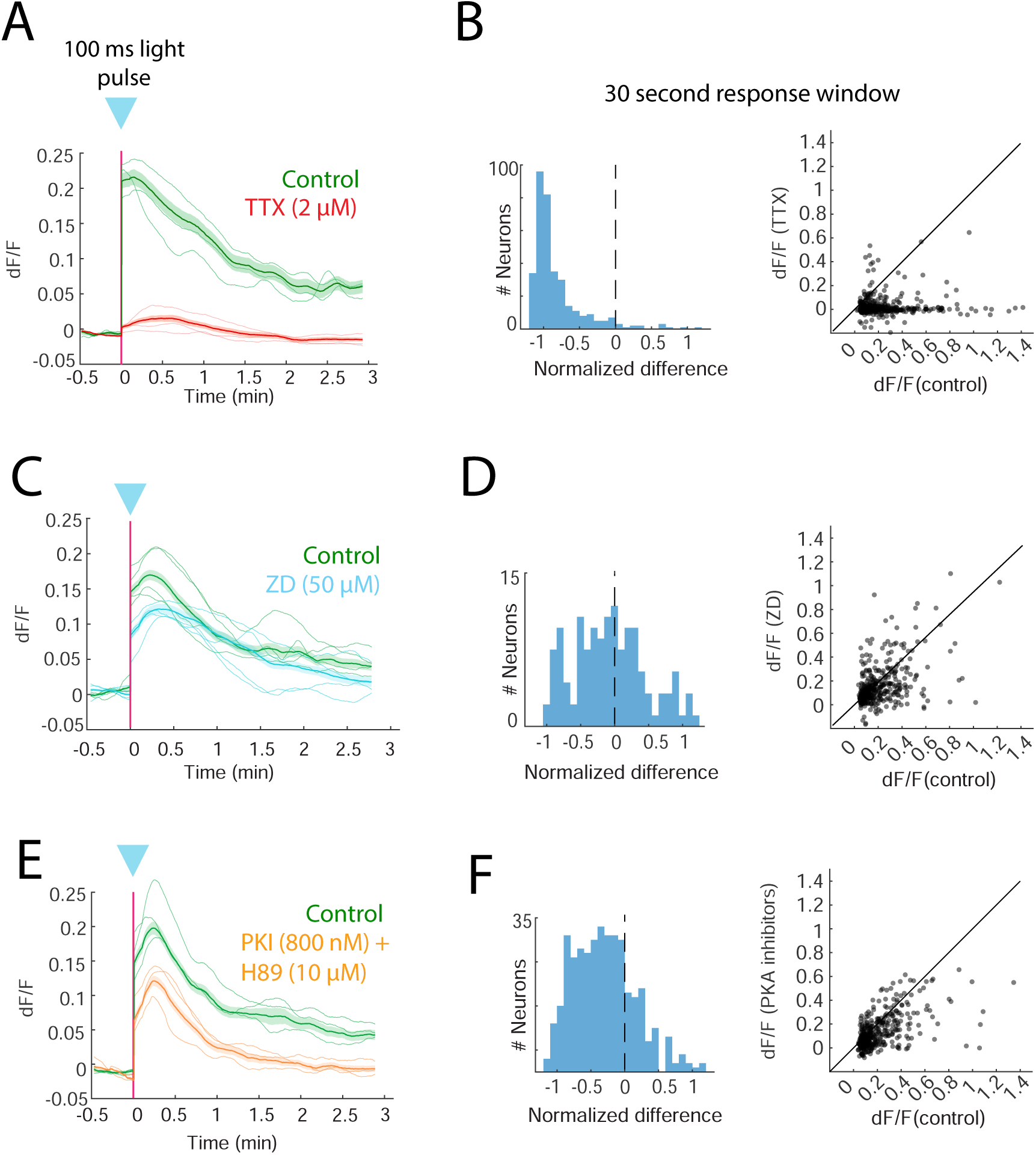
Effect of TTX, ZD, and PKA inhibitors on cAMP-evoked neural activity. Related to Figure 4. A) Average GCaMP responses to blue light stimulation of biPAC before and after TTX application (2 μM, 3 slices from 2 mice). B) Histogram showing the relative change ((drug - baseline) / baseline) of early (averaged over the first 20 seconds after stimulation) dF/F responses to blue light stimulation before and after TTX application. Right: scatter plot of dF/F responses before and after TTX application (20 μM). Mixed-effects model, two-sided permutation test. TTX effect size: −0.23 dF/F. ***p < 0.00. (283 responsive neurons, 3 slices, 2 mice). C,D) Similar to A-B, but for application of the HCN channel blocker ZD7288 (50 μM). Mixed-effects model, two-sided permutation test. ZD effect size: −0.045 dF/F. ***p < 0.001. (383 responsive neurons, 4 slices, 3 mice). E,F) Similar to A-B, but for application of the PKA blockers PKI (800 nM) and H89 (10 μM). Mixed-effects model, two-sided permutation test. Drug effect size: −0.075 dF/F. ***p < 0.001. (411 responsive neurons, 4 slices, 3 mice).

## Methods

All animal care and experimental procedures were approved by the Beth Israel Deaconess Medical Center Institutional Animal Care and Use Committee. Animals were singly housed on a 12-h light/dark cycle with standard mouse chow and water provided ad libitum unless they were used for feeding-related experiments, in which case they were food restricted (~2.5 g of standard chow per day for a maximum of 2 weeks, with daily weight monitoring) to maintain body weight at 10 – 15% of free-feeding weight. All experiments were performed on adult Vglut2-ires-cre mice (Jackson Lab Strain # 016963; 29 male and 17 female mice in total) between the ages of 8 and 20 weeks.

### Viral injections

Injections (~150 nl total) were targeted to the ventral edge of PBN (AP: −5.05 mm; DV: −3.8 to −3.7 mm; ML: ± 1.5 mm) to allow for spread of virus via efflux. For experiments with simultaneous biPAC stimulation and cADDis recordings, we injected a 1:1 mixture of AAV1-hSyn-DIO-GreenDownward-cADDis (Boston Children’s Hospital Vector Core) and AAV1-EF1a-DIO-mKate2-biPAC (Boston Children’s Hospital Vector Core). For cell-attached experiments, we injected a 1:1 mixture of AAV1-EF1a-DIO-Venus-biPAC and AAV1-EF1a-DIO-mKate2-PDE4D3-cat (Boston Children’s Hospital Vector Core), or a 1:1 mixture of AAV1-EF1a-DIO-Venus-biPAC and phosphate buffered saline. For optogenetic stimulation experiments, we used AAV1-EF1a-DIO-mKate2-biPAC followed by implantation of optic fibers. For photometry recordings of PKA activity, we injected AAV2/9.hSyn.FLEX.ExRai-AKAR2-CW3SL (gift from Huganir lab) followed by implantation of optic fibers. For experiments testing the role of cAMP signaling on feeding behavior, we either injected mice with AAV1-EF1a-DIO-mKate2-biPAC or AAV1-EF1a-DIO-mKate2-PDE4D3-cat.

### Fiber photometry recording

Optic fibers with a metal ferrule (400 μm diameter core; multimode; numerical aperture (NA) 0.48; 5.0 mm length; Doric Fibers; MFC_400/430-0.48_5mm_MF1.25_FLT) were bilaterally implanted over PBN (Bregma: AP: −5.05 mm; DV: −3.65 mm; ML: ±1.5 mm). The fibers and a custom-made titanium headpost (H.E. Parmer) were fixed to the skull using C&B Metabond (Parkell). Mice were given at least 2 weeks to recover before any behavioral experiments began. Fiber photometry recordings were performed using head-fixed mice that were free to run on a 3D printed circular treadmill. Fiber optic cables (1 m long; 400 μm core; 0.48 NA; low autofluorescence; Doric Lenses) were coupled to implanted optic fibers with zirconia sleeves (Precision Fiber Products). Black heat shrink material was placed around the fiber coupling to prevent light leakage from interfering with recordings or behavior. Excitation and emission light was passed through a GFP fluorescence minicube (FMC3_E(460–490)_F(500–550); Doric Lenses). Excitation light (~100 μW) was provided by a 465 nm LED (Plexon LED and driver) which was modulated at either 217 or 319 Hz using transistor–transistor logic (TTL) output from two lock-in amplifiers (SR830; Stanford Instruments). Emission light was collected by a femtowatt photoreceiver (Newport 2151), demodulated using a lock-in amplifier (SR830; Stanford Instruments) and digitized at 1 kHz sampling rate (PCIe-6321; National Instruments). Data acquisition was controlled using a custom script in MATLAB (MathWorks).

### GRIN lens implantation and related surgical procedures

For PBN cell body imaging, immediately after viral injections, mice were implanted with a gradient index (GRIN) lens. We used singlet GRIN lenses (3 mice, GRINtech, NEM-050–25–10–860-S-1.5p; 0.5 mm diameter; 6.5 mm length; 250 μm focal distance on brain side at 860 nm (NA 0.5); 100 μm focal distance on air side (NA 0.5); non-coated) and doublet GRIN lenses (5 mice, GRINtech, NEM-050-25-10-860-DM; 0.5 mm diameter; 9.89 mm length; 250 μm focal distance on brain side at 860 nm (NA 0.47); 100 μm focal distance on air side (NA 0.19)). GRIN lens implantations were performed as previously described^1^. Briefly, mice were maintained under anesthesia (1.5–2.0% isoflurane) and body temperature was maintained using a heating pad in a stereotaxic apparatus (Kopf Instruments, Model 940). A beveled 25-gauge needle (0.51 mm diameter, Fisher) was attached to the stereotaxic holder and zeroed on Bregma. To give the lens a snug fit and reduce brain motion artifacts, the needle was inserted slowly to a depth of 0.1 mm higher than the final depth of the GRIN lens. After this needle was removed from the brain, the GRIN lens was held with a bulldog serrafine clamp (Fine Science Tools, catalog no. 18050–28) with heat shrink on the tips (to improve grip and prevent damage) and zeroed at Bregma without touching the skull (to avoid debris covering the lens surface). The lens was then slowly inserted in the hole made by the needle and down to its final depth, then secured to the skull by applying Metabond (Parkell) or ultraviolet-curable glue (Loctite) around the lens. A titanium headplate was centered over the GRIN lens and fixed to the skull using Metabond. Afterwards, a 3D-printed plastic funnel was cemented onto the headplate, which allowed the securing of a light shield between the head and the objective using Velcro, to prevent collection of stray light. After completion of the GRIN lens surgery, the top of the GRIN lens was protected by a cut-off tip of an Eppendorf tube (Fisher) and secured using Kwik-Cast (WPI). The mice were allowed to recover from surgery for at least 2 weeks before any behavioral experiments.

### Cell-attached recordings

Brain slices were prepared by deeply anesthetizing animals (6-8 weeks old; ad libitum fed mice) with isoflurane followed by rapid decapitation. Upon removal, brains were immediately immersed in ice-cold, carbogen-saturated (95% O2, 5% CO2) choline-based cutting solution consisting of (in mM): 92 choline chloride, 10 HEPES, 2.5 KCl, 1.25 NaH2PO4, 30 NaHCO3, 25 glucose, 10 MgSO4, 0.5 CaCl2, 2 thiourea, 5 sodium ascorbate, 3 sodium pyruvate, oxygenated with 95% O2/5% CO2, measured osmolarity 310 – 320 mOsm/L, pH= 7.4. Then, 275-300 µm-thick coronal sections containing the PBN complex were cut with a vibratome (Campden 7000smz-2) and incubated in oxygenated cutting solution at 34°C for 10 min. Next, slices were transferred to oxygenated aCSF (126 mM NaCl, 21.4 mM NaHCO3, 2.5 mM KCl, 1.2 mM NaH2PO4, 1.2 mM MgCl2, 2.4 mM CaCl2, 10 mM glucose) at 34°C for an additional 15 min. Slices were then kept at room temperature (20–24°C) for ~45 min until use. A single slice was placed in the recording chamber where it was continuously superfused with oxygenated aCSF at a rate of 3-4 mL per min at room temperature. Neurons were visualized with an upright microscope (Olympus) equipped with infrared-differential interference contrast and fluorescence optics. All recordings were made using a Multiclamp 700B amplifier and data were filtered at 2 kHz and digitized at 20 kHz. Loose-seal, cell-attached recordings (seal resistance 20-50 MW) were made in voltage-clamp mode with the recording pipette filled with aCSF and holding current maintained at V_h_ = 0 mV. To photoactivate biPAC, a LED light source (470 nm) was used. The blue light was focused onto the back aperture of the microscope objective (40X) producing wide-field exposure around the recorded cell of 10-15 mW per mm^2^ as measured using an optical power meter (PM100D, Thorlabs). A programmable pulse stimulator, Master-8 (A.M.P.I.) and pClamp 10.2 software (Molecular Devices, Axon Instruments) controlled the photostimulation output.

### Freely moving feeding experiments with optogenetic stimulation

Mice were injected with biPAC bilaterally and a titanium headpost was cemented onto their skull. After 4 weeks of expression, mice were habituated to being connected to patch cords (bilateral, 400μm core, 0.39NA; Doric lenses) in their home cages (2 days; 1 - 2 hours per day). 24 hours before a test session, all food and bedding were removed from the cage and replaced with fresh bedding without food. During the test session, these mice were connected to patch cords in their home cage and provided with a single ~3 g standard chow pellet that was weighed beforehand. The pellet was then weighed every 30 minutes for 2 hours during which time the mice remained either unstimulated or received photostimulation either for 90 minutes following a 30 minute unstimulated period (100 ms pulse every 10 seconds; 1 mW 473 nm laser; LaserGlow) or during the first 30 minutes only (1 ms pulse every 10 seconds; 5 mW 473 nm laser; LaserGlow).

### Head-fixed feeding experiments with optogenetic stimulation or tail shocks

We used a head-fixed behavioral paradigm^1,2^ to achieve a higher temporal resolution assessment of the influence of photostimulation on feeding. Following freely moving photostimulation experiments, the same mice were habituated to head-fixation for 2 days. Mice were then maintained on a food restricted diet and trained to lick at a lick spout to receive Ensure. Following 2 days of habituation to the lick spout, mice were allowed to consume Ensure freely for 20 minutes. On test days with photostimulation, mice received 5 photostimulation trains (1 s on, 2 s off, 10 second duration, 1 mW, 465 nm LED; Plexon; two minute inter-train interval).

For experiments testing the effect of tail shock on feeding behavior, a separate cohort of mice were prepared with headpost implants only. Mice were chronically food restricted and trained to lick to receive Ensure as described above. On test days, mice received a 10 second duration tail shock train (6 Hz; 0.35 mA) that began 2 minutes after onset of free binging on Ensure. During initial experiments, we repeated this tail shock stimulus 5 times (2 minute intervals). In most experiments after the initial 5 mice, we only used a single tail shock train per trial and allowed several minutes breaks between trials to ensure the presence of multiple trials with stable licking before a tail shock (typically 2 - 3 trials per day).

To test the role of cAMP signaling on the duration of tail shock-evoked suppression of feeding, we performed unilateral viral injections of a cre-dependent AAV-PDE4D3-Cat, a constitutively active phosphodiesterase that should selectively disrupt cAMP only^4^. We chose to use unilateral injections as we found that bilateral expression of PDE4D3-Cat in PBN neurons led to high mortality of the mice. Following five weeks of expression, mice were tested on the licking behavior as described above. We compared these animals to a set of control mice that expressed hM4Di-mCherry in PBN neurons but that were not injected with CNO. Posthoc histological analysis showed no obvious cell death resulting from expression of PDE4D3-Cat.

### Analysis of head-fixed licking behavior

For statistical analysis of the influence of manipulations (photostimulation or tail shock trains) on ongoing binge licking, we compared licking during the 1 minute (for photostimulation experiments) or 2 minutes (tail shock experiments) following the manipulation vs. the same window of time in unstimulated control sessions.

### Two-photon imaging via GRIN lenses

Two-photon imaging was performed using a resonant-scanning two-photon microscope (Neurolabware) at 15.5 frames s^−1^ and 796 × 512 pixels per frame as described previously^1,5^. Excitation was achieved using an InSight X3 laser (Spectra-Phsyics). Imaging was performed with a 10x 0.5 NA air objective (TL10X-2P; ThorLabs). Imaging fields of view were at a depth of 100–300 μm below the face of the GRIN lens and generally targeted the dorsal lateral PBN (4 mice) or external lateral PBN (3 mice; ad libitum fed). We imaged using an excitation wavelength of 960 nm for all GCaMP or cADDis imaging, and 1100 nm for measuring biPAC fluorescence as a proxy for protein expression. Laser power ranged between 40 and 60 mW at the front aperture of the objective (the power at the sample was substantially less because of partial transmission via the GRIN lens).

### Two-photon imaging of acute brain slices

For two-photon imaging of acute brain slices, slices were prepared as described above and transferred to a recording chamber perfused with ACSF (oxygenated with 95% O2 and 5% CO2; flow rate: 2–5 mL/min) at room temperature. All mice used for slice experiments were ad libitum fed. Imaging was performed with a 16×0.8 NA water-immersion objective (Nikon). The excitation wavelength used was 920 nm. In optogenetic experiments involving biPAC, the PMT was powered off during the photostimulation (100 ms to 2 s; 470 nm LED; 1 mW/mm^2^, Luxeon Star LEDs), which was driven by an Arduino-controlled driver (Luxeon Star LEDs).

For all pharmacological manipulations except kynurenic acid application, we collected pre- and post-drug responses to biPAC stimulation in the same cells. To block sodium channels, we applied tetrodotoxin (TTX, Sigma Aldrich, 2 uM in ACSF). To block PKA activity we used a mixture of myristoylated protein kinase A inhibitor (PKI, 800 nm) and H89 (10 mM). To block hyperpolarization-activated cyclic nucleotide-gated channels, we used the selective blocker ZD7288 (50 mM, Tocris). Finally, to block excitatory transmission, we incubated and recorded slices in ACSF containing kynurenic acid (KA, Sigma Aldrich, 200 nM). All drugs were applied for at least 15 minutes before starting a new biPAC stimulation trial.

### Two-photon *in vitro* and *in vivo* image processing

Image registration for all two-photon calcium imaging of cell bodies was performed using Suite2p^6^. For extraction of signals from cell body regions of interest (ROIs) from volumetric brain slice imaging (15 depths; 10 um apart), we used CellPose^7^, which optimally identified ROIs from GCaMP- or cADDis-expressing cell bodies from the mean image of the movement-corrected field of view. Fluorescence time series were extracted by averaging each of the pixels within each binarized mask. We calculated neuropil activity as the median value of an annulus surrounding each ROI (inner radius: 15 pixels; outer radius: 50 pixels; pixels belonging to any other ROI were excluded from these annulus masks).

### Two-photon data analysis

This time course of neuropil activity was subtracted from the activity time course of the associated ROI to create a fluorescence time course, F(t), where t is the time of each imaging frame. The change in fluorescence was calculated by subtracting a running estimate of baseline fluorescence (F0(t)) from F(t), then dividing by F0(t): dF/F(t) = (F(t) - F0(t))/F0(t), where F0(t) is the average fluorescence in a 30 second window prior to biPAC stimulation, or a 5-minute window prior to drug application.

Significantly responding neurons (in the case of both cADDis and GCaMP) were defined as neurons where average post-stimulation fluorescence in the first 20 seconds after the blue light pulse exceeded baseline by 1 standard deviation and 0.05 dF/F. Of note, cADDis is a downwards sensor (i.e. it signals through fluorescence decreases), and so we have flipped all cADDis traces and signals to be positive for ease of visualization throughout the paper.

To calculate the duration of GCaMP responses to biPAC stimulation, we calculated the amount of time that each neuron’s GCaMP fluorescence exceeded a pre-determined threshold (exceeding 0.05 dF/F and exceeding the fluorescence level during the 30 second baseline period by at least 1 standard deviation) until the first time it returned below this threshold for at least 20 consecutive seconds. Neurons were classified as transient if their GCaMP response lasted for less than 60 seconds, and as sustained if the response lasted more than 60 seconds.

To quantify the rate of clearance of cAMP in each neuron, we used the following formula:

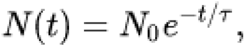

Where (N_0_) is the starting cADDis level (80% of peak cADDis fluorescence), N(t) is the time at which fluorescence reaches 20% of peak, λ is the decay constant, computed by fitting a monoexponential spanning N_0_ to N(t), and τ is the decay time constant. Of note, the temporal dynamics of our GCaMP and cADDis responses to biPAC stimulation were slower *ex vivo* than observed in our *in vivo* experiments (Figure 3D; median *ex vivo* cADDis duration above threshold, 68 ±12.3 s; median *τ,* 70.6 s; Figure 4D, median *ex vivo* GCaMP duration above threshold, 48.9 ±1.5 s), potentially due to the difference in temperature (25° C *ex vivo*).

### Statistics

No statistical methods were used to pre-determine sample sizes, but our sample sizes were chosen to reliably measure experimental parameters while remaining in compliance with ethical guidelines for minimizing animal use and were similar to those reported in previous publications. Experiments were conducted by an investigator with knowledge of the animal genotype and treatment. All of the virus expression, optic fiber implants and GRIN lens placements were verified by post hoc histology or were observable during the experiment (slice imaging). All data presented as bar and line graphs indicate mean ± s.e.m., with individual data points also plotted. Parametric statistical tests were used in most cases. In some, linear mixed effect models were used (see below). Statistical analyses were performed in Matlab (Mathworks). Significance levels are indicated as follows unless otherwise specified: \**p*< 0.05; \*\**p*< 0.01; \*\*\**p*< 0.001.

### Mixed-effects models

Linear mixed effects models were used to account for the hierarchical dependencies in the data (neurons −> slices −> mice), particularly when a large sample of neurons was being compared across conditions. Specifically, we accounted for correlations between measurements that were sampled from the same slice by including random effect terms for slice ID and for mouse ID. Fixed effect coefficient and P values are reported in the main text and figures.

